# Artificial Selection of Multispecies Microbial Inoculants

**DOI:** 10.64898/2026.11.17.767241

**Authors:** Belen Benitez-Dominguez, Ignacio Belda, Alvaro Sanchez

## Abstract

Evolutionary engineering can optimize microbial consortia through iterative ecological perturbation and selection. It remains unclear whether selection improves community function simply by enriching communities with the strongest individual contributors or whether it can also enrich communities with synergistic interactions among species. Here, we address this question by optimizing the initial composition of multispecies wine yeast inoculants. Each generation we independently perturb the starting abundance of each of six yeast strains, toggling it between two of six possible doses (from 0 to 10^7^ CFU/mL), and select for two functions of contrasting complexity. Unlike previous studies focused on serially passaged communities, our approach optimizes the initial inoculum composition for single-batch fermentation, a mode of operation common to many biotechnological processes. Selecting for a function that all individual strains can independently contribute to results in inoculants that are enriched with the strongest contributors. In contrast, selecting for a function that no strain can perform alone yields more complex communities whose performance emerges from synergistic interactions. These findings show that evolutionary engineering can exploit both individual contributions and interspecies interactions to optimize microbial communities. Our approach can be adapted to optimize other microbial inoculants of low-to-medium diversity.

## Introduction

Microorganisms inhabit virtually every environment in the biosphere, and humans have long harnessed their metabolic capabilities, from their traditional use in food production to more recent roles in biofuel production and drug manufacturing [1]. Although modern biotechnology has largely relied on engineered microbial monocultures, growing recognition of the ecological contexts in which microbes naturally occur has shifted attention toward the biotechnological potential of microbial communities [2–4]. By distributing metabolic tasks among specialists, microbial communities can perform complex functions while reducing the burden on individual members, while offering greater robustness and resilience across environmental conditions [5]. Thus, biotechnology has recently expanded its focus from optimising a single strain to engineering entire communities.

Evolutionary engineering methods, such as artificial community-level selection, represent a promising approach to designing microbial communities. Artificial selection offers a path to iteratively explore consortium compositions to optimise a target function: one starts by generating a library of variant communities, keeps those that perform best and uses them as the basis for a new library of variants.

This process is repeated over successive rounds to improve the function of interest. These methods have been used to tune the growth environment using genetic algorithms [6, 7] and to select whole communities by their collective function, propagating the best performers across rounds [8–14]

To date, artificial community-level selection has been applied to serially passaged microbial communities [15–18]. However, many processes of biotechnological interest, such as food fermentations, operate in a single batch mode [19]. In single batch culture, one starts by seeding the initial inoculant into the bioreactor. After that initial inoculation, species grow and interact, often exhibiting substantial ecological turnover and large changes in population size. As this ecological succession unfolds, a function of interest is produced, and the process ends when the product is collected. A classic example of a single batch biotechnological process is wine fermentation. During wine fermentation, ecological processes, such as environmental filtering, nutrient competition and rising ethanol toxicity, drive a pronounced succession from a diverse initial consortium that includes many non-*Saccharomyces* yeasts to the dominance of *Saccharomyces cerevisiae* [20]. The initial composition can therefore influence both the course of fermentation and the properties of the finished wine. Consequently, wine making relies heavily on optimising the starting inoculants. Because strains that contribute during the early stages of fermentation may be rare or absent by its end, the community recovered at the end of one fermentation does not necessarily contain the strains that mattered at its start, and cannot simply be carried over as the next inoculum. The challenge is therefore to select the initial inoculum composition that generates a desirable fermentation trajectory. The composition of an inoculant is defined not only by which species are present, but by the density or population size at which each species is introduced, and both jointly determine the succession and its endpoint. Because all species can be included from the outset, what remains to be designed is a continuous quantity, i.e., the set of starting population sizes across all species. This defines a space far too large to empirically characterize exhaustively, calling for an evolutionary engineering approach.

In this paper, we ask whether evolutionary engineering can optimise the composition of multispecies inoculants for two target functions of significance for wine fermentation: maximising sugar consumption in a challenging high-sugar, fructose-rich must, and producing lactic acid in fermentations that also finish sugar consumption. Because these two functions are very different in terms of their presumed complexity, they allow us to address an unresolved question in community-level artificial selection [18, 21]: is a positive response to selection primarily caused by an enrichment of strains that perform well on their own or by selecting species combinations with strong synergistic interactions?

As a model system to tackle these questions, we used an inoculant consisting of six different yeast strains (three *Saccharomyces cerevisiae*, and three non-*Saccharomyces*), which were inoculated at different cell densities. Over three rounds of selection, we fermented 64 candidate inocula, kept the best-performing one, and used it to define the next 64 by varying the initial cell density of each strain around its winning value. For both targets, we found an inoculant that improved function over the best inoculant in the starting generation and also outperformed every single-species inoculant. The basis of improvement differed depending on the function being optimised. Selection for sugar consumption, a function every strain can contribute to on their own, improved it by tuning the doses of highly-contributing individual strains. In contrast, selection for the combined requirement of finishing fermentation and producing lactic acid, which no single species could achieve alone, favored both individual strain contributions and strong pairwise interactions. Thus, our results provide a blueprint for optimising microbial inoculants for biotechnologically relevant applications and show how individual strain contributions and species interactions can contribute differently to improvement depending on the inherent complexity of the function being selected.

## Results

The purpose of our experiments is to apply evolutionary engineering to search the design space of all possible inoculants one may form from six yeast strains (Fig. 1A): three strains of *S. cerevisiae* (SC1, SC2, SC3), and one each of *Torulaspora delbrueckii* (Td10), *Metschnikowia pulcherrima* (Mp21) and *Lachancea thermotolerans* (Lt24). All six strains are present in every inoculant, so what distinguishes one inoculant from another is not which species are included but the population size at which each is inoculated. We explored a range spanning five orders of magnitude, from 10³ to 10^7^ CFU/mL, plus the absence of a strain as the lowest level. Allowing each of the six strains six possible doses in that range defines 6^6^ = 46,656 candidate inoculants.

**Fig 1.**
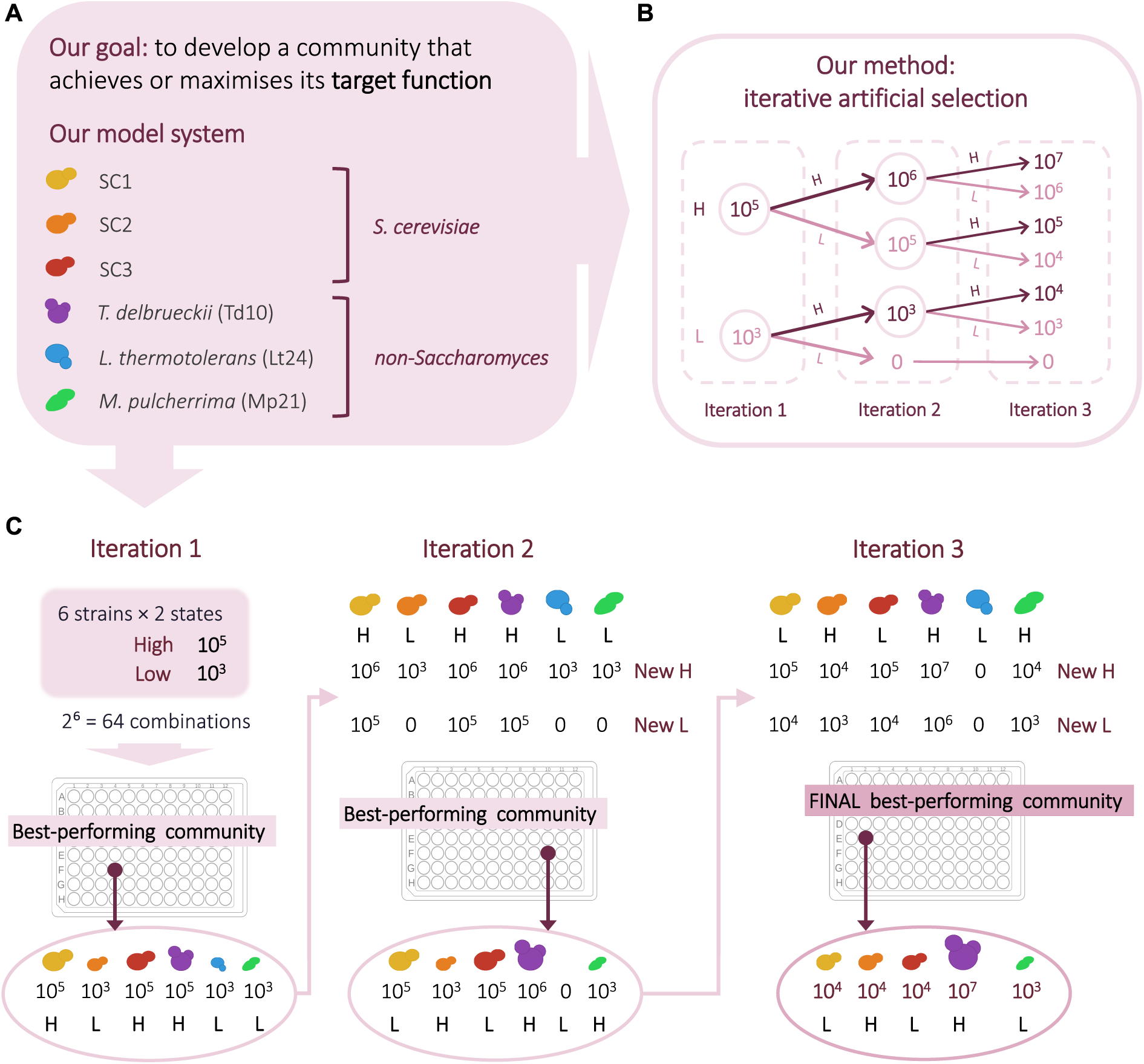
Iterative artificial selection of inoculum doses. **(A)** From a fixed pool of six commercial wine yeasts, three *Saccharomyces cerevisiae* (SC1, SC2, SC3) and three non-*Saccharomyces* (*Torulaspora delbrueckii*, Td10; *Lachancea thermotolerans*, Lt24; *Metschnikowia pulcherrima*, Mp21), we select the inoculant that best performs a target function. **(B)** Each strain is set at each round to one of two doses, low (L) or high (H). The best inoculant sets the dose window for the next round: for each strain, its winning dose becomes one bound of a refined window, shifted up or down according to whether it won high or low (initial levels 10³ and 10^5^ CFU/mL; ceiling 10^7^ CFU/mL). **(C)** In each round the six strains toggled between their two current doses give 2^6^ = 64 candidate inoculants, assembled from pure cultures, fermented and assayed for the function; the best-performing community defines the doses explored in the next round. Because the inoculant is rebuilt from pure cultures each round, what is selected is the set of doses, not a propagated community, and an optimised consortium is reached within three rounds.

In each round, every strain is toggled between two doses, a low and a high level, so that only 2^6^ = 64 inocula need to be assayed (Fig 1B). In the first round these levels were 10³ and 10^5^ CFU/mL for every strain. We fermented the 64 communities and measured the target function (Methods). To create the next generation, we followed a simple hill-climbing scheme. Rather than propagating the winning community, we kept the single best-performing inoculum (Fig 1C) and redefined the two doses of each strain around its winning dose, shifting the window up or down according to whether it had won at its high or low level (Fig 1B; Methods). The 64 inocula of the next generation were assembled from pure cultures, and three rounds were run for each function.

### Two target functions at contrasting levels of functional complexity

We selected for two functions that differ in functional complexity, by which we mean whether each strain can contribute to the function on its own, or whether the function can only be reached through interactions between strains (Fig 2). We reasoned that selection on a simple function would favour inoculants carrying high doses of the strongest individual contributors, whereas selection on a complex function would favour inoculants with strongly interacting strains, since no strain can perform the function alone.

**Fig 2.**
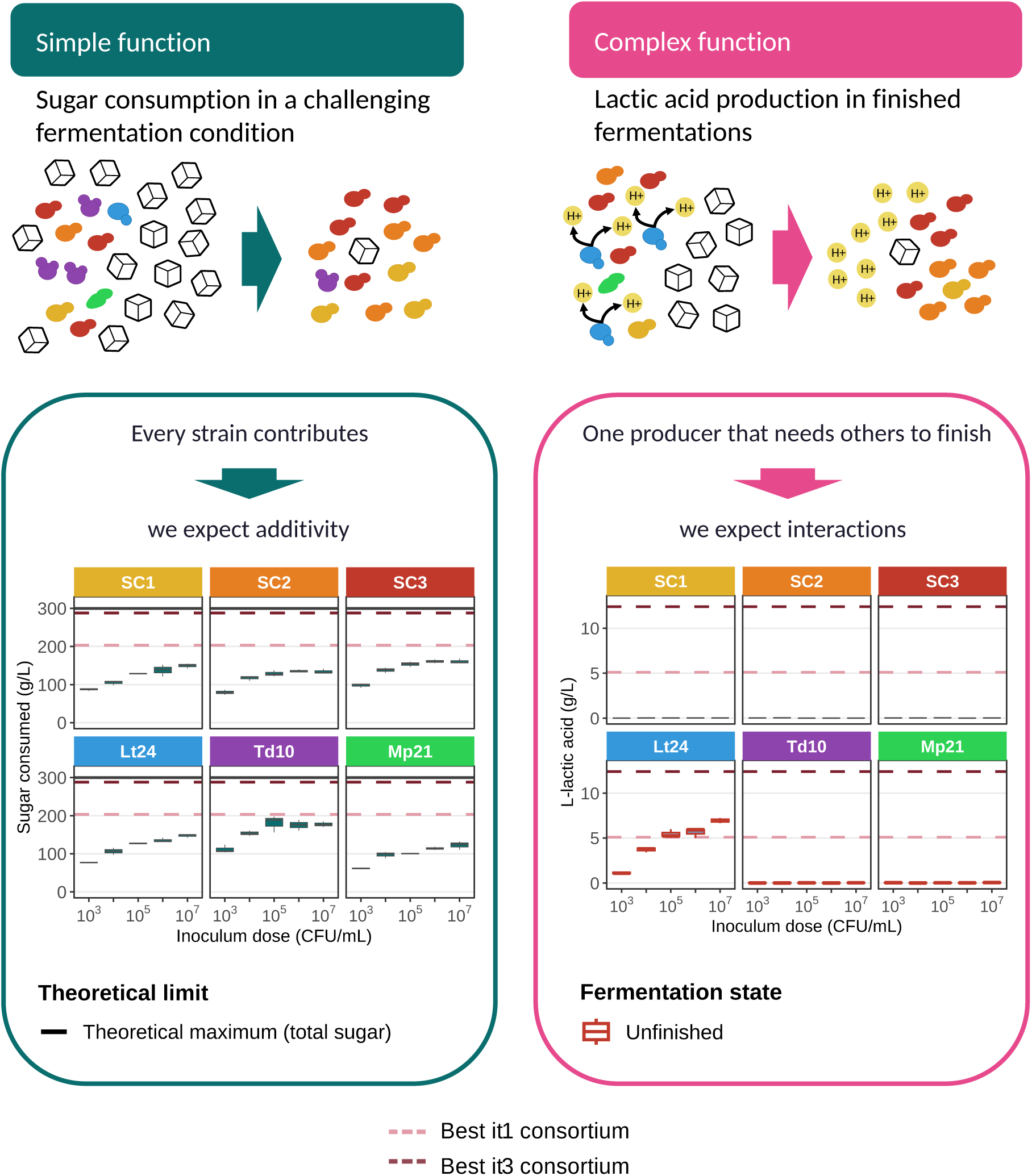
Two target functions at contrasting levels of functional complexity, and the monoculture performance that motivates selecting communities. The optimised consortium from the third round of selection (Fig 1) is characterised for two functions chosen to differ in how the task maps onto the pool. **Left, simple function: sugar consumption**. The assay is a must (**HS-FR** must) that makes fermentation harder than a standard synthetic grape must (SGM), carrying more sugar (300 g/L instead of 200 g/L) and a higher proportion of fructose (a 2:1 fructose:glucose ratio instead of 1:1); osmotic stress and the excess of fructose make it difficult for the fermentation to finish. A fermentation was considered finished when no more than 5 g/L of sugar remained. *Lower panel:* sugar consumed by each strain in monoculture across five inoculation doses. Solid line, theoretical maximum (total sugar available). **Right, complex function:** L-lactic acid production, assayed in **SGM** and scored only in communities that finished the fermentation. *Lower panel:* L-lactic acid produced by each strain in monoculture across the same doses; red outlines mark fermentations that did not finish. In both right and left lower panels, plots show three replicates per dose (inoculation dose 10³ to 10^7^ CFU/mL); dashed lines mark the function reached by the best consortium of the first (pink) and third (dark red) rounds of selection.

As the simple function we chose sugar consumption, assayed in a high-sugar, fructose-rich must (HS-FR) that carries 300 g/L of sugar, rather than the 200 g/L of a standard synthetic grape must (SGM), at a 2:1 fructose-to-glucose ratio instead of the usual 1:1 (Fig 2, left). These conditions are sub-optimal for *S. cerevisiae*. The high sugar concentration raises the osmotic pressure of the must, which slows growth and fermentation and makes sluggish or stuck fermentations more likely, a problem that is becoming more common as warmer climates raise grape sugar at harvest [22]. And because *S. cerevisiae* is glucophilic, taking up fructose less efficiently than it does glucose [23], the later, fructose-dominated stages ferment slowly, the more so as ethanol accumulates and further impairs fructose uptake. We scored the function as the total sugar consumed on day 7. Because every strain in the pool consumes sugar, even though none finishes the fermentation on its own (Fig 2, left), improvement can draw on all six, and we expected it to rest mostly on additive contributions.

As the complex function we chose L-lactic acid production, assayed in standard SGM and scored only in communities that finished fermentation, that is, with no more than 5 g/L of residual sugar (Fig 2, right). L-lactic acid lowers the pH of the wine through biological acidification, which is useful where musts from warm regions lack acidity [24, 25], but only once the fermentation has finished, as industrial practice requires. We therefore scored it as a step function, counting the acid a community produced only when it finished the fermentation and setting the function to zero otherwise, regardless of the acid accumulated (formalised in Methods). Among the pool, only *L. thermotolerans* produces L-lactic acid, by diverting part of the glycolytic pyruvate to lactate during alcoholic fermentation [24], yet it cannot finish the fermentation at any dose, while the three *S. cerevisiae* strains finish the fermentation but produce no acid (Fig 2). Because producing the acid and finishing the fermentation fall to different strains, and the step function rewards a community only when both are achieved, a high function value cannot be reached by any strain alone and requires pairing a producer with a finisher. We therefore expected selection to rely on a synergy between strains, imposed by how we define the function.

### Sugar consumption in high-sugar fructose-rich musts improves through additive strain effects and is dominated by a single species

Our first experiment selected for maximal sugar consumption in the HS-FR must, which contains 300 g/L of sugar in a 2:1 fructose:glucose ratio. All six strains in our pool consume sugar under our growth conditions, although none ferments it completely in monoculture (Fig 2, left). For all three rounds of inoculants we determined the whole distribution of the function across the 64 inocula (Fig 3A), and the function of the best consortium improved by 40.6% over three rounds of selection (Fig 3B). In the first round the distribution was clearly bimodal, splitting into a lowand a high-consuming group according to whether *Torulaspora delbrueckii* was set at its low (10³ CFU/mL) or high (10^5^ CFU/mL) level (Fig 3A). Before any model was fitted, the shape of the distribution already pointed to this strain as decisive.

**Fig 3.**
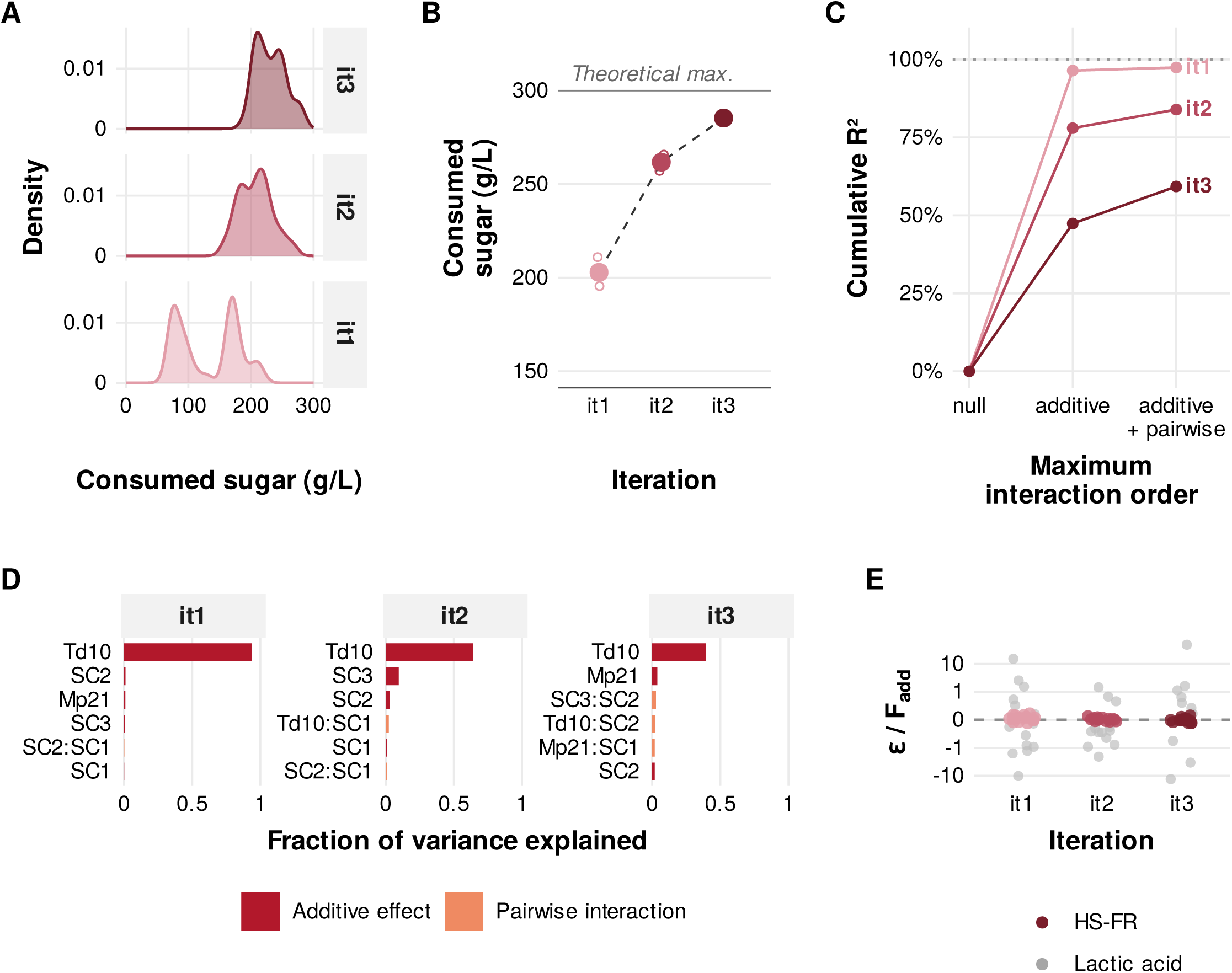
Sugar consumption in the HS-FR must is largely additive and dominated by a single strain. Directed evolution of the inoculum in a high-sugar (300 g/L), 2:1 fructose:glucose must, across three rounds (it1–it3). **(A)** Distribution of sugar consumed across the 64 communities at each round. **(B)** Trajectory of the winning consortium, reaching a total gain of +40.6%; filled circles are the mean and open circles the three replicates. The solid line marks the theoretical maximum (300 g/L). The winning inocula were: it1, Mp21 10^5^, Lt24 10^5^, Td10 10^5^, SC3 10^5^, SC2 10³, SC1 10³; it2, Mp21 10^5^, Lt24 10^5^, Td10 10^6^, SC3 10^6^, SC2 10³, SC1 10³; it3, Mp21 10^4^, Lt24 10^4^, Td10 10^6^, SC3 10^7^, SC2 10^4^, SC1 10^4^ (CFU/mL). **(C)** Cumulative variance explained (R²) as terms up to a given interaction order are added. **(D)** The six terms contributing the most to the variance in each round, with single-strain and pairwise terms coloured differently. **(E)** Size of each pairwise interaction relative to the additive expectation, *ε*/F_add (Methods), for every pair and round, from the additive-plus-pairwise model. HS-FR is coloured by round; lactic acid (Fig 4) is shown in grey for comparison. The y axis is pseudo-logarithmic (linear near zero, logarithmic for large values) so that both signs and zero can be shown. Colours denote round in panels A–C and E.

To determine whether the improvement came from stronger individual contributions or from interactions between strains, we partitioned the variation in function across the 64 communities at each generation. We coded each strain by whether it was inoculated at its low or high dose and fitted two nested models, one in which strains act independently and their effects simply add up, and one that also allows the effect of each strain to depend on the dose of another (pairwise interactions). The gain in variance explained when interaction terms are added measures how much of the function rests on interactions rather than on independent strain effects (Methods). Individual contributions accounted for most of the explained variation, with the additive model reaching R² = 0.96, 0.78 and 0.47 across the three rounds (Fig 3C), and *T. delbrueckii* was the largest single contributor in every round (Fig 3D). Pairwise terms added 1%, 6% and 12% of the variance. They did not exceed what experimental noise alone produces in the first two rounds, but did in the third (S1 Fig), as communities approached the optimum. The interactions that did appear were also small relative to the function itself. Expressed as a fraction of the value expected if the two strains simply added their effects, no pairwise interaction departed from that expectation by more than 11%, and the median departure was 2–4% in each round (Fig 3E). The variance left unexplained by the two models grew across rounds. Selection therefore improved the function mainly by increasing the doses of the strongest individual contributors, rather than by exploiting interactions

### Optimal lactic acid production in wine inocula requires interactions between complementary strains

The second experiment involved a more complex function, one that, as argued above, was more likely to involve interactions. In our monocultures, *L. thermotolerans* was the only strain that produced L-lactic acid; yet, on its own it does not finish the wine fermentation at any dose tested (Fig 2, right). By contrast, the three *S. cerevisiae* strains are able to finish fermentation on their own (i.e. no more than 5 g/L sugar is left), yet they produce no lactic acid. Selection was therefore constrained on two fronts at once: only communities that finished fermentations were scored, and among these we quantified lactic acid as described in the Methods and selected the top community. After applying the same evolutionary algorithm as in the previous experiment, we found that the function of the best community in each round responded positively to selection. Over three rounds the best consortium raised its lactic acid titre by 134%, and, just as in the previous experiment, the whole distribution of inocula shifted towards higher mean functions (Fig 4A, B). In the first round that distribution had a marked peak near zero, made up of communities that finished the fermentation but produced little or no acid, those carrying *L. thermotolerans* at its low dose; as selection raised the producer’s dose, this peak gave way and the bulk of communities moved to higher titres (Fig 4A).

**Fig 4.**
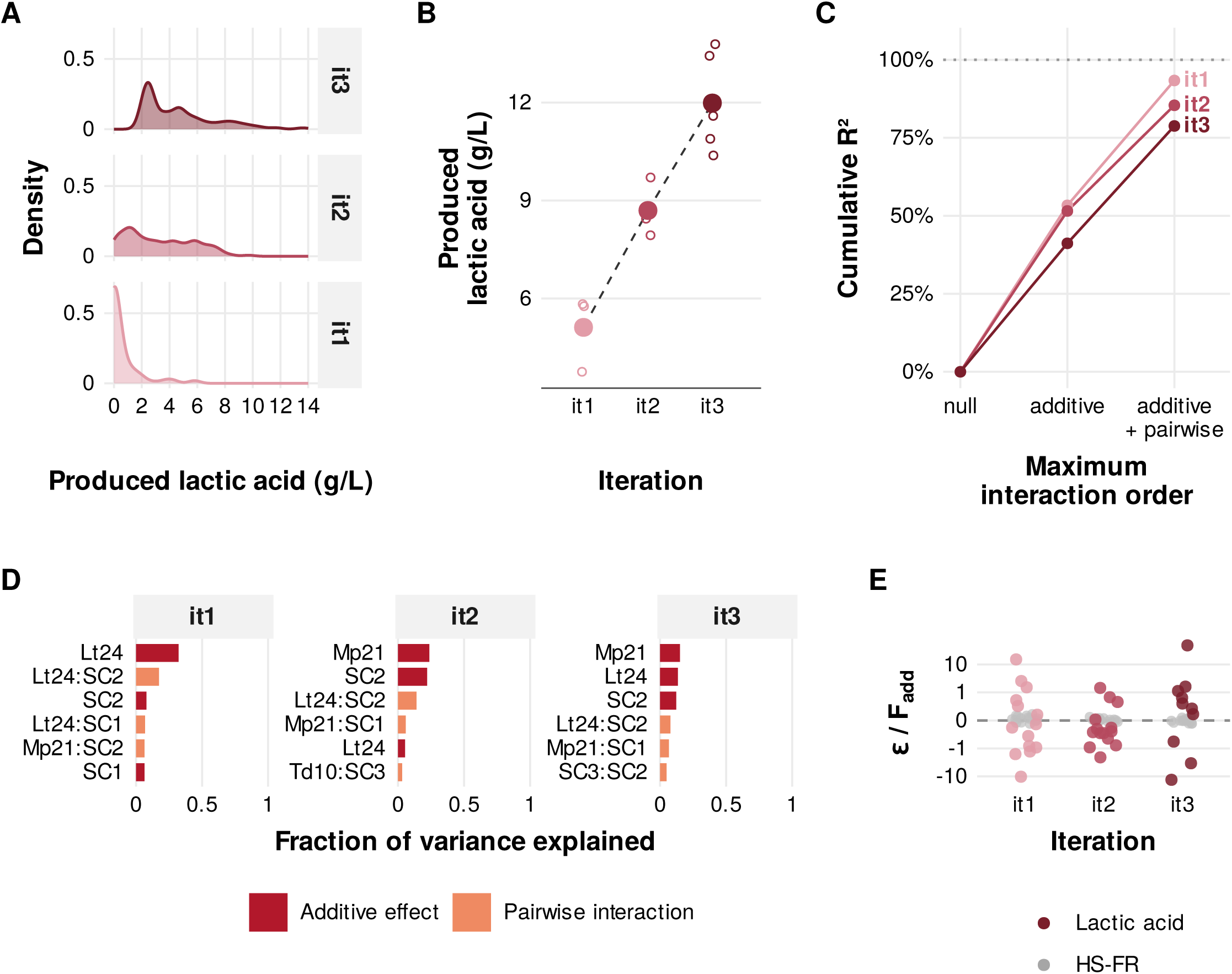
Lactic acid production is interactive. Directed evolution of the inoculum for L-lactic acid production in synthetic grape must across three rounds (it1–it3). The function was scored only in communities that finished the fermentation (no more than 5 g/L of residual sugar) and set to zero otherwise. **(A)** Distribution of lactic acid produced across the 64 communities at each round. **(B)** Lactic acid produced by the winning consortium in each round. Filled points give the mean and open circles the technical replicates. The winning inocula were: it1, Mp21 10³, Lt24 10^5^, Td10 10³, SC3 10^5^, SC2 10³, SC1 10³; it2, Mp21 10³, Lt24 10^6^, Td10 absent, SC3 10^6^, SC2 10³, SC1 10³; it3, Mp21 10^4^, Lt24 10^7^, Td10 absent, SC3 10^7^, SC2 10^4^, SC1 10^4^ (CFU/mL). **(C)** Cumulative variance explained (R²) by the null, additive and additive-plus-pairwise models, fitted to the communities that finished the fermentation. **(D)** The six terms contributing the most to the variance in each round, with single-strain and pairwise terms coloured differently. **(E)** Strength of each pairwise interaction relative to the additive expectation, *ε*/F_add (Methods), per strain pair and round, for lactic acid (colour) and for sugar consumption in the HS-FR must (grey; Fig 3E). The y axis is pseudo-logarithmic (linear near zero, logarithmic for large values) so that both signs and zero can be shown. Colours denote round in panels A–C and E.

Notably, the optimised consortium produced more L-lactic acid than *L. thermotolerans* did at any dose in monoculture (Fig 2). The partner strains therefore did not merely allow the fermentation to finish but raised the producer’s output above its own monoculture level. Consistent with our original expectations, selection for this more complex function produced communities enriched in interactions. Although each strain’s independent contribution explained similar share of the variance, pairwise terms explained 38% of it in the third round, the only one with all 64 communities (Fig 4C), well above what experimental noise alone produces (S1 Fig), whereas in sugar consumption they exceeded that noise only in the third round. Interactions were also large relative to the function itself. The median pairwise interaction amounted to 34–88% of the value expected from additive effects, and in some pairs it exceeded that value (Fig 4E). As we expected, the strongest interactions involved *L. thermotolerans* and the *S. cerevisiae* strains (Fig 4D).

### Landscape ruggedness explains why selection enriched communities with strong interactions

At the outset of our experiments we had hypothesised that Lt24 would interact synergistically with the *Saccharomyces* strains, as it is the only one to produce lactic acid while the *Saccharomyces* are necessary to finish the fermentation. Our regression analysis supports a role for interactions. Pairwise terms carried a substantial share of the variance in every round (38% in the complete third round), and the producer was involved in the strongest of them (Fig 4D). The single largest pairwise term, Lt24×SC2, is present in all three rounds. The interaction structure is also richer than a single producer-finisher pair. Further terms appear, among them an interaction between *Saccharomyces* strains (SC3×SC2) and terms involving Mp21. This indicates that the effect of each strain on the function depends strongly on the doses of the other strains present. As we show below, this dependence can reverse the sign of a strain’s effect (Fig 5B). Such sign reversals make the community-function landscape we explore through artificial selection rugged, that is, able to hold more than one peak, so that raising each strain towards its individually best dose does not necessarily lead to the best community [26, 27]. This would explain why artificial selection enriched interactions for this function, rather than simply increasing the inoculum sizes of the strains that contribute most on average, as it did for sugar consumption.

**Fig 5.**
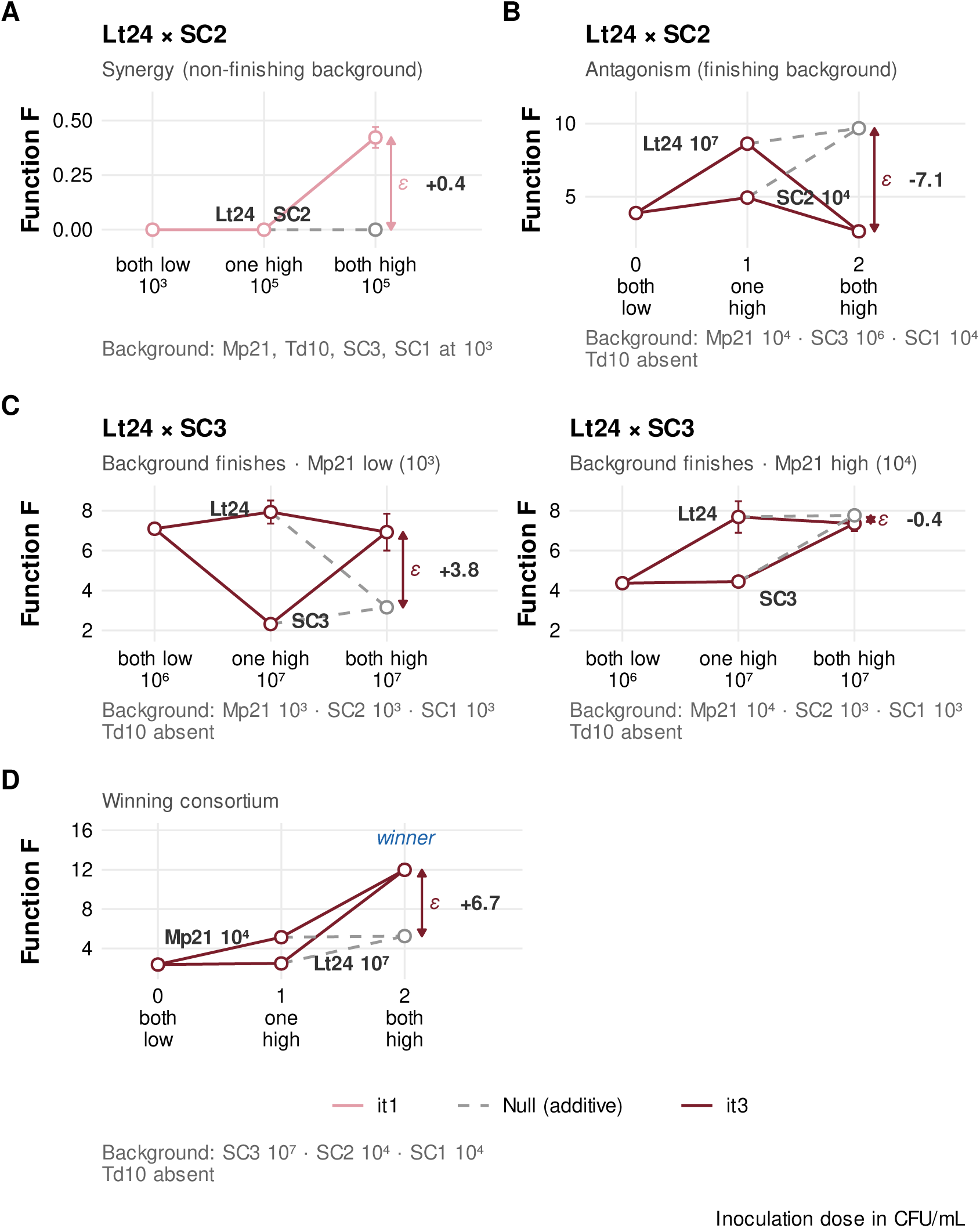
Pairwise interactions are context-dependent. Epistasis diamonds for pairs involving the producer Lt24. The x axis gives how many strains of the pair are at their high dose, with the doses toggled given on the axis or next to each strain; the four remaining strains are fixed at the background shown below each panel. Nodes are communities (mean of the two plate positions of the same combination in the third round) and error bars are the standard error of the technical replicates. The grey dashed line and open circle are the additive expectation, and *ε* is the measured both-high node minus that expectation. **(A)** Lt24 × SC2 in the first round, in a background that does not finish the fermentation, so the function stays at zero unless both strains are raised (*ε* = +0.4), a positive interaction imposed by the finishing requirement rather than by metabolism. **(B)** The same pair in the third round, in a background that finishes the fermentation; raising SC2 together with the producer lowers the acid below what either reaches alone (*ε* = *−*7.1). **(C)** Lt24 × SC3 in the third round with Mp21 at its low (left) or high (right) dose; the positive interaction at low Mp21 (*ε* = +3.8) disappears when Mp21 is raised (*ε* = *−*0.4). **(D)** Mp21 × Lt24 within the winning lactic consortium (*ε* = +6.7).

To gain a finer understanding of the context-dependence of species effects and the ruggedness of the landscape, we adopted the epistatic view of species interactions, in which an interaction is present when a species modifies the functional effect of another species in a given community background [28–35]. This view resolves interactions one background at a time, which a regression term, being an average over backgrounds, cannot do. The strongest interaction identified by the regression, Lt24×SC2, changes sign between rounds. In the first round, where finishing the fermentation is the limiting step, it is positive (Fig 5A). This positive interaction is imposed by the way the function is scored rather than by metabolism. Lactic acid is produced only when the producer is high, and the fermentation finishes only when a finisher is high, so the function rises above zero only when SC2 and the producer are both raised. By the third round every community finishes fermentation, and there the same pair shows reciprocal sign epistasis (Fig 5B). Each strain raises the acid on its own, but raising SC2 together with the producer lowers it below what either reaches alone, even though the fermentation still finishes, so SC2 suppresses production rather than failing to finish. The interaction between two strains can also depend on a third. The interaction between Lt24 and SC3, for instance, is positive when Mp21 is at its low dose and disappears when Mp21 is raised (Fig 5C).

Because interactions are contextual, what matters for optimisation is not how two strains interact on average but how they interact inside the communities selection actually reached. In the winning lactic consortium the two non-Saccharomyces are required together. With the three *Saccharomyces* at their higher dose, neither Mp21 nor Lt24 reaches the optimum when raised on its own, and only raising both does (12 g/L, *ε* = +6.7 ; Fig 5D). The consortium that selection produced for this function is therefore enriched in interactions of the kind described here. Optimising over a rugged landscape by directed evolution yielded a community whose function emerges from interactions between strains, rather than from an enrichment of the strains that contribute most additively, as selection did for the simpler sugarconsumption function.

## Discussion

Six strains at six possible doses define 46,656 candidate inocula, a space too large to assemble and measure exhaustively. Directed evolution of the inoculum reached an optimised consortium for each of the two functions after measuring 64 inocula in each of three rounds, and in every case the whole distribution shifted upward from round to round, not only the best community. Two features made this work. First, selection is directed, so each round is informed by the last and a landscape of this size need not be surveyed in order to be navigated. Second, the strains are varied by dose rather than by inclusion, whereas bottom-up assembly and composition-function landscapes have been mapped almost entirely over the presence or absence of each member [30, 34]. In tuning how much of each strain to add rather than whether to include it, our scheme is the ecological counterpart of moving from a gene knockout to gene dosage. That this dimension matters is clear from the first round in the HS-FR must, where inocula of identical membership split into two performance groups by the dose of *T. delbrueckii* (Td10) alone (Fig 3A).

We selected for two functions chosen to differ in how the task maps onto the pool, and the response to selection followed that difference. Sugar consumption, a function every strain can contribute to, behaved as expected. Pairwise terms explained 1% and 6% of the variance in the first two rounds, within what experimental noise alone produces, and 12% in the third, the only round in which they exceeded that null (S1 Fig). Even then, no interaction departed from the additive expectation by more than 11% (Fig 3E), so improvement rested on single-strain effects. Lactic acid production, which only one strain can perform and only in a fermentation that others finish, was more interactive. Pairwise terms explained 38% of the variance in the complete third round, against 12% for sugar consumption (Figs 3C and 4C), and individual interactions were often comparable in size to the value expected from additive effects (Fig 4E). The same six strains thus produced two very different interaction structures, set not by the pool but by the function being selected.

This bears on a common assumption about consortia, that distributing a function across several strains entails metabolic interactions between them, as in the cross-feeding and metabolic interdependencies through which division of labour is usually framed [36, 37]. Both functions are distributed in this sense, in that no monoculture reached, at any dose we tested, the function attained by the optimised consortium (Fig 2). For lactic acid the reason is specific. The strain that produces most acid alone, *L. thermotolerans* (Lt24), does not finish the fermentation, and only finished fermentations were scored. In sugar consumption, by contrast, every strain consumes sugar on its own, so sharing the task creates no dependencies between strains and the function remains mostly additive. Whether a consortium can be designed one strain at a time therefore depends on whether its strains depend on each other to perform the function, not on how many of them contribute to it.

In the HS-FR must, sugar consumption is dominated by a single strain, *T. delbrueckii*, whose contribution is large and, across most of the landscape, independent of the rest of the inoculum, so its effect adds on rather than depending on which other strains are present. The optimum, however, sits where that independence breaks down. In a community of *S. cerevisiae* that already consumes most of the sugar, raising *T. delbrueckii* from 10^6^ to 10^7^ CFU/mL (the two doses tested in that round) reduces the function instead of adding to it (from 285 to 210 g/L; S2 Fig). Lactic acid production is the opposite case from the start, since it requires two things at once that no strain delivers together. *L. thermotolerans* makes the acid but fades as ethanol rises and cannot finish the fermentation, while the S. cerevisiae strains finish it but do not make the acid.

The two optima are shaped by interactions in opposite directions. The lactic optimum is built on cooperation. With the fermentation finished by the *S. cerevisiae*, the function still requires the two non-*Saccharomyces* together, and neither the producer nor *M. pulcherrima* (Mp21) reaches the optimum when raised on its own (Fig 5D). The producer also realises more of its potential inside a finishing community than in monoculture, because the strains that finish the fermentation let it keep producing (Fig 2). The high-sugar optimum is reached the other way, by keeping an interference out. Because *T. delbrueckii* interferes once the *S. cerevisiae* are in place, selection keeps it at the lower of its two doses rather than raising it. Improving a function was therefore as much about which strains to hold down as about which to raise.

*T. delbrueckii* shows why a strain cannot be ranked on its own. It is the largest single contributor to sugar consumption (Fig 3D), yet the best consortium holds it down in both functions, at the lower of its two doses in the high-sugar must and out of the inoculum altogether for lactic acid (Fig 4B). What a strain does is therefore set by the community it is in and not by the strain alone.

Read this way, the additivity of a function follows from its biology. A function dominated by one strain acting monotonically and independently tends to add up, whereas one that hinges on a threshold or on conflicting requirements between strains does not. Additivity is not even a fixed property of a whole function. Sugar consumption is additive across most of its landscape and yet interactive near its optimum, the region selection has to reach, which is where pairwise terms first exceeded experimental noise (S1 Fig). Statistical models built on single-strain effects recover most of this landscape [30, 34, 38], yet they misfit its extremes, and the maximum is one of them. Because the additive fit credits *T. delbrueckii* with its positive average effect, it ranks the inoculum carrying the strain at its highest dose as the best, whereas, as described above, raising it to that dose in a community that already completes the consumption lowers the function, so the additively best choice is not the best consortium. Our results suggest that whether additive or interactive effects matter is better assessed function by function, and background by background, than assumed for a community as a whole.

None of this is particular to wine. We tuned inocula rather than propagating whole communities because the ecology of these systems rules propagation out. Many microbial processes run as single batches with strong ecological turnover [19], in which an added inoculum shapes the early community but is overtaken as succession proceeds, so the community that performs the function is not necessarily one that can be carried over to the next round. Wherever this holds, the inoculum is effectively the only thing that can be set in advance, and our results show that designing it round by round can improve a target function quickly, while revealing how much of that function depends on individual strains and how much on the way they are combined.

## Materials and Methods

### Yeast strains and inoculum preparation

We used a fixed pool of six selected wine yeasts: three *Saccharomyces cerevisiae* strains (SC1, SC2 and SC3) and three non-*Saccharomyces* strains, one each of *Torulaspora delbrueckii* (Td10), *Metschnikowia pulcherrima* (Mp21) and *Lachancea thermotolerans* (Lt24), all provided by Lallemand Bio. These are commercial products, and we identify them hereafter by these codes only: SC1, SC2, SC3, Td10, Mp21, and Lt24, respectively. We started precultures from single colonies on plates prepared the day before and grew them in 50 mL of Yeast Extract Peptone Dextrose (YEPD) in 100 mL Erlenmeyer flasks for 24 h at 25 °C with shaking at 200 min^-1^. We then harvested the cells and washed them four times in PBS by centrifugation (2500 rpm, 25 min), removing residual YEPD nutrients, before adjusting cell density in the corresponding synthetic grape must and using them as inoculum in experimental fermentation trials. Communities differed in the dose of each strain rather than in which strains were included, except where a strain’s lower dose had fallen to zero.

### Fermentation conditions and functions

We ran fermentations statically in deep-well 96-well plates, each well holding 600 µL of must under a gas-permeable lid, at 25 °C for seven days, and then measured the target function. A fermentation was considered finished when no more than 5 g/L of sugar remained.

We assayed two functions, each in a dedicated synthetic grape must (SGM) prepared as in Ruiz et al. [33]:(i) a high-sugar, fructose-rich must (HS-FR), an SGM modified to contain 300 g/L of sugar (against the usual 200 g/L) in a 2:1 fructose-to-glucose ratio (200 g/L fructose and 100 g/L glucose, rather than an equal split). We measured the function as total sugar consumed on day 7; (ii) standard SGM (200 g/L of sugar; with a 1:1 glucose:fructose ratio; as prepared in [33]) for L-lactic acid production. Because acidification only matters in a finished wine, we scored this function only in communities that finished fermentation (*≤*5 g/L residual sugar on day 7), as the L-lactic acid produced. We define the scored function as F = L · H(c *−* s), where L is the L-lactic acid concentration, s the residual sugar on day 7, c = 5 g/L the finishing threshold, and H the Heaviside step function (H = 1 when the fermentation finished, s *≤* c; H = 0 otherwise). A fermentation that did not finish therefore contributes F = 0 by construction, regardless of the lactate it accumulated.

We quantified reducing sugars by the dinitrosalicylic acid (DNS) method. Samples were diluted 1:20, mixed with DNS reagent (10 g/L 3,5-dinitrosalicylic acid, 16 g/L NaOH, 30 g/L potassium sodium tartrate) in a 96-well plate, incubated at 65 °C for 1 h, cooled to room temperature, and read at 540 nm against a standard curve run in parallel.

L-lactic acid was quantified enzymatically in microplate with an L-lactate dehydrogenase (LDH) coupled colorimetric assay, in which L-lactate oxidation reduces NAD^+^ to NADH and NADH reduces nitroblue tetrazolium to a formazan via phenazine methosulfate. Samples were diluted 1:10 and combined with LDH, NAD^+^, phenazine methosulfate and nitroblue tetrazolium in TRIS-HCl buffer (0.7 M, pH 9), incubated at 28 °C for 30 min in the dark, developed with DMSO, and read at 570 nm against a standard curve run in parallel.

### Directed evolution of inocula

We optimised each function over successive rounds of selection. In every round we assigned each of the six strains one of two inoculation doses, a low (L) and a high (H) level, giving 2^6^ = 64 candidate inocula that span the corners of the dosage space. We assembled the 64 communities in parallel from pure cultures, fermented and measured them as above, and identified the best-performing inoculum. In the first round of the HS-FR experiment, the two inocula with the highest mean consumption differed by 0.4 g/L (203.3 and 202.9 g/L), far less than the variation between their replicate wells (standard deviations of 25.1 and 7.8 g/L). We took as the winner the one whose replicates were more consistent. Rather than propagating this community, we used it to redefine the dose window for the next round, refining the dose of each strain around the value it took in the winner. In the first round the two levels were 10³ and 10^5^ CFU/mL. The same rule applied in later rounds around the best performer, shifting a strain’s window downward when it kept winning low (holding at 0 once reached) and upward when it kept winning high, up to a ceiling of 10^7^ CFU/mL, beyond which higher doses have little biological meaning. We ran three rounds of selection for each function, by which point the strains that kept winning high had reached this ceiling in both experiments.

In the third round of the lactic acid experiment both dose levels of *T. delbrueckii* were 0, so each of the 32 dose combinations of the remaining five strains occupied two positions of the plate layout, giving six replicate wells per combination instead of three. The community at position H04, whose inoculum had been prepared incorrectly, was fermented again and its new values were used.

### Variance decomposition by interaction order

For each function and round, we represented the inoculum of each community by a binary vector **x** = (*x*_1_, . . ., *x_N_*) *∈* {0, 1}*^N^*, where *x_i_* = 0 and *x_i_* = 1 denote the low and high inoculation doses of strain *i* within that round’s dose window. Because all strains are present in every community, *x_i_* encodes dose rather than presence, except where the low dose of a strain had fallen to zero. The community–function landscape is the function *F*(**x**), sugar consumption or lactic acid production, averaged across replicate wells. Since the dose window is redefined every round, each round was analysed separately and all quantities describe the landscape locally, within that round.

All three HS-FR rounds comprised the 2^6^ = 64 dose combinations. In the lactic acid condition only communities that finished the fermentation were scored, which left 33 and 44 of the 64 combinations in the first and second rounds. In the third round *T. delbrueckii* was absent from every inoculum, so its variable was constant and was removed (*N* = 5), and the 64 communities covered each of the 32 combinations of the remaining strains twice. Both communities of each combination entered the analysis as separate observations.

We fitted an additive model and an additive-plus-pairwise model by ordinary least squares,

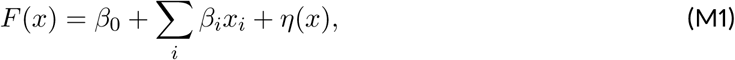

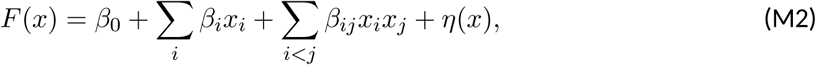

where *η*(**x**) is the residual error. With this coding, *β*_0_ is the function with all strains at their low dose, *β_i_*the effect of raising strain *i* to its high dose with all other strains low, and *β_ij_*the change in that effect when strain *j* is also at its high dose. We quantified the variance explained by each model *m* with the coefficient of determination

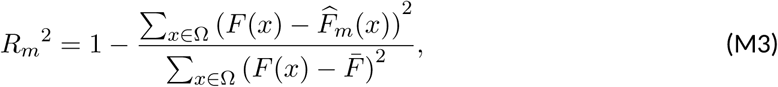

where Ω is the set of observed combinations, *F*^^^*m* the prediction of model *m* and *F* the mean observed function. *R*²_1_ is the variance explained by single strains, *R*²_2_ *− R*²_1_ the variance added by pairwise terms, and 1 *− R*²_2_ the variance left unresolved, which combines interactions among three or more strains and measurement noise (Figs 3C and 4C). Terms beyond pairs were not fitted because they cannot be estimated in the incomplete rounds, and the same two models were fitted in every round so that rounds and functions could be compared.

### Distinguishing pairwise interactions from experimental noise

Measurement noise alone makes the additive-plus-pairwise model explain more variance than the additive model. To test whether pairwise terms explained more than this, we simulated 1000 landscapes without interactions for each round,

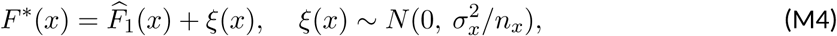

where *σ_x_* is the standard deviation across the *n_x_* replicate wells of community **x**. We fitted both models to each simulated landscape and considered the observed increase *R*²_2_ *− R*²_1_ significant when it exceeded the 95th percentile of the simulated increases (S1 Fig).

### Variance explained by individual terms

To identify which strains and pairs contributed most to the variance in each round (Figs 3D and 4D), we computed the sum of squares explained by each term *S* of the model of Eq. (M2), entering single-strain terms first, and expressed it as a fraction of the total,

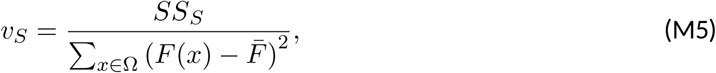

where *S* is a single strain or a pair. The fractions of single-strain terms add up to *R*²_1_ and those of pairwise terms to *R*²_2_ *− R*²_1_. Figs 3D and 4D show the six terms with the largest *v_S_*in each round. In complete rounds *v_S_* does not depend on the order in which terms are entered, whereas in the incomplete first and second lactic acid rounds it does, so their partition is approximate.

### Communities without lactic acid

In the first lactic acid round, 18 of the 33 communities that completed fermentation produced no acid. All 15 producing communities contained *L. thermotolerans* at its high dose, whereas all 18 non-producing communities contained it at its low dose. Thus, the presence or absence of lactic acid was fully determined by the dose of this strain among the observed communities, although this does not rule out interactions affecting the amount produced. From the second round onwards, every community that finished fermentation produced lactic acid.

### Background-dependent pairwise interactions

The coefficient *β_ij_*of Eq. (M2) describes the interaction between strains *i* and *j* with all other strains at their low dose, as estimated by the model. To examine how this interaction depends on the doses of the other strains, we measured it directly in specific backgrounds ***b***, defined by the doses of the *N −* 2 other strains,

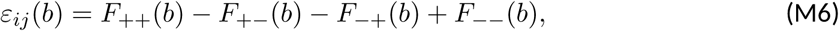

where *F_pq_*(***b***) is the measured function with strains *i* and *j* at doses *p* and *q*, respectively, + denoting the high and *−* the low dose. *ε_ij_*(***b***) is the departure of the community with both strains at their high dose from the value expected if their separate effects added up, positive for synergy and negative for antagonism. A dependence of *ε_ij_*(***b***) on ***b*** means that other strains modify the interaction, which is the signature of higher-order interactions. For these comparisons the lactic acid function was set to zero in communities that did not finish the fermentation, so that all 64 communities could be used. Thus, in the first two lactic acid rounds, these interactions were calculated from a different landscape than the regression coefficients. In the third lactic acid round each node combines the replicate wells of both positions of that combination. Nodes in Fig 5 and S2 Fig show the mean of the replicate wells and error bars the standard error.

To compare the size of interactions across pairs, rounds and functions (Figs 3E and 4E), we normalised the interaction predicted by the second-order model (Eq. M2) by its additive expectation, evaluated in a fixed background with all other strains at their low dose,

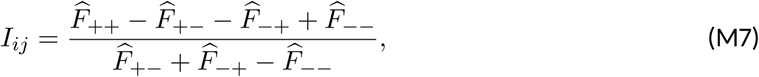

All *F*^^^ values in Eq. (M7) were calculated from the fitted model of Eq. (M2), with the remaining strains at their low dose. With the 0/1 coding this ratio equals *β_ij_*/ (*β*_0_ + *β_i_*+ *β_j_*). Pairs whose interactions could not be estimated from the available data were excluded.

### Comparison with Walsh–Hadamard decomposition

In the complete rounds (the three HS-FR rounds and the third lactic acid round), we also decomposed the variance with the Walsh–Hadamard expansion, which includes terms of every order (S1 Methods) [28]. Single-strain and pairwise terms explained fractions of the variance similar to those of the regression models. Terms involving three or more strains exceeded experimental noise in the third round of both functions, as expected when the function approaches saturation.

### Software

All analyses were carried out in R 4.4.2 [39]. Linear models were fitted with the stats package, and figures were produced with ggplot2 4.0.3 [40] and patchwork 1.3.0 [41]. Code and data are available at this repository

## Supporting information

Supplementary informatioin

## Acknowledgements

The authors acknowledge Lallemand Oenology for providing the strains and funding this project through contracts with UCM and IBFG-CSIC (title: Optimising the performance of wine yeast consortia through rational combinatorial design). The fundamental research that enabled this work was supported by MI-CIU/AEI/10.13039/501100011033 and the European Regional Development Fund (ERDF; "A way of making Europe"), through grants supporting the laboratories of Alvaro Sanchez (PID2021-125478NA-I00) and Ignacio Belda (PID2022-138343NB-I00). Belén Benítez-Domínguez acknowledges her predoctoral grant PRE2022-103063 funded by MICIU/AEI/10.13039/501100011033.

We thank Sergio Izquierdo, Isabel Quirós and Andrea Arrabal for their help with the experimental work, Jose Camacho-Mateu for theoretical and data analysis help in relation with the Walsh-Hadamard transform, Alba Rodriguez for administrative assistance, as well as all other members of of the Sánchez laboratory for their feedback on the paper and valuable discussions throughout this project.

