## Supplementary informatioin for "Artificial Selection of Multispecies Microbial Inoculants"

#### Supporting information

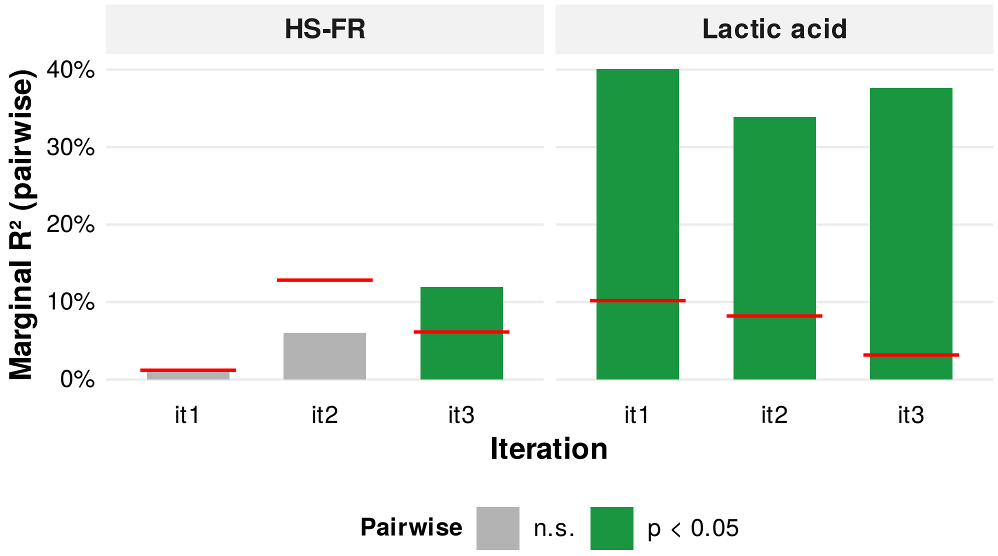


**S1 Fig. Contribution of pairwise interactions compared with experimental noise.** For each function and round, the bar gives the increase in the variance explained, $R^2$, when pairwise terms are added to the additive model, $R^2{}_2 - R^2{}_1$ (Methods). The red line marks the 95th percentile of the same increase in 1000 simulated landscapes without interactions, in which the function was generated from the fitted additive model plus Gaussian noise scaled to the measured error between replicate wells of each community. Bars above the line (green) indicate that pairwise terms explain more variance than noise alone would produce ($p < 0.05$), and bars below it (grey) that they do not. Pairwise terms did not exceed the null in the first two HS-FR rounds and exceeded it in the third HS-FR round and in all three lactic acid rounds.

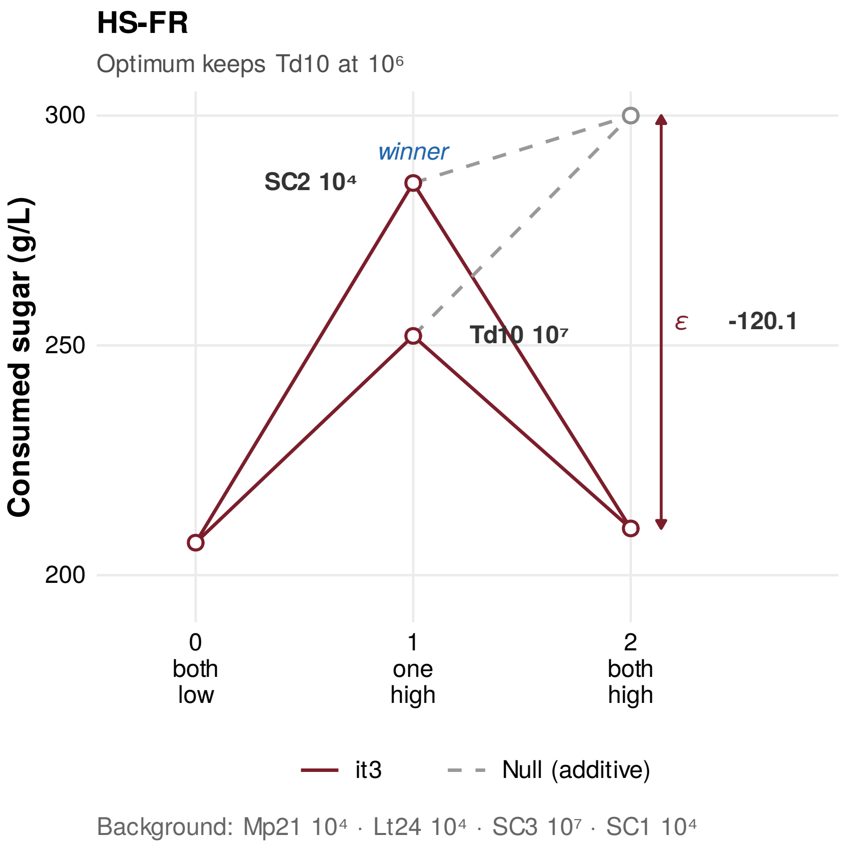


**S2 Fig. Interaction between *T. delbrueckii* (Td10) and *S. cerevisiae* SC2 near the high-sugar optimum (third round).** Epistasis diamond for the two strains in the winning HS-FR background, with *M. pulcherrima* and *L. thermotolerans* at their low dose and *S. cerevisiae* SC3 and SC1 at their high dose. Each node is one measured community, and its error bar is the standard error over three replicates. The function runs from both strains at their low dose, through each one at its high dose alone, to both at their high dose, and $\varepsilon$ is the departure of the both-high node from the sum of the two separate effects, negative for antagonism. The additive expectation for both strains at their high dose exceeds the sugar available, about 300 g/L, and is capped at that value in the figure, whereas $\varepsilon$ is computed from the uncapped expectation. Taking 300 g/L as the expectation instead, $\varepsilon$ would be $-90$ g/L.

528 **S1 Methods.** Walsh–Hadamard decomposition of the community–function landscapes. Includes Fig A.

### S1 Methods. Walsh–Hadamard decomposition of the community–function landscapes

We represented inoculum composition by a binary vector $\mathbf{x} = (x_1, \cdots, x_N) \in \{-1, 1\}^N$, where $x_i = -1$ and $+1$ denote the low and high inoculation doses of strain $i$, respectively. We used $-1/+1$ rather than the $0/1$ coding used in the main text, following the convention of the Walsh–Hadamard expansion introduced below. The community-function landscape is a function $F(\mathbf{x})$ assigning a quantitative output to each inoculum composition: sugar consumption or lactic acid production, averaged across replicate wells. In the third lactic acid round each combination occupied two positions of the plate layout, and the wells of both positions were pooled as replicates of that combination (six wells).

We applied the expansion only to the rounds in which every dose combination was measured, the three HS-FR rounds (N = 6, $2^6$ = 64 combinations) and the third lactic acid round (N = 5, $2^5$ = 32 combinations, with *T. delbrueckii* absent). The first two lactic acid rounds, with 33 and 44 of the 64 combinations, were analysed only with the regression models of the main Methods. Since the dose window is redefined every round, all quantities describe the landscape locally, within each round.

For completely sampled landscapes, containing all $2^N$ inoculum compositions $\mathbf{x}$, we used the Walsh–Hadamard (WH) expansion to decompose community function into additive, pairwise, and higher-order terms

$$F(\mathbf{x}) = \beta_0 + \sum_i \beta_i x_i + \sum_{i<j} \beta_{ij} x_i x_j + \sum_{i<j<k} \beta_{ijk} x_i x_j x_k \cdots . \quad (1)$$

#### Variance explained by each interaction order

The WH decomposition Eq. (1) provides an exact orthogonal decomposition of the total function variance

$$V_{tot} = \langle F^2 \rangle_{\mathbf{x}} - \langle F \rangle^2{}_{\mathbf{x}} = \sum_{k=1}^{N} V(k) . \quad (2)$$

The order-resolved variance associated with interaction order $k$ is

$$V(k) \; = \sum_{|\,S|\,=k} {\beta_S}^2, \quad (3)$$

where $S \subset \{1, 2, \, ... \, , N\}$ denotes a subset of strains, $|S| = k$ is the number of strains in that subset, and $\beta_S$ is the corresponding WH coefficient. We quantified the relative contribution of each interaction order as $V(k)/V_{tot}$.

To assess whether the variance associated with each interaction order exceeded experimental noise, we built 1000 noise-only landscapes following Camacho-Mateu et al. [28]. For each combination, the deviations of its replicate wells from their mean were scaled by $\sqrt{}$(n/(n − 1)), so that they kept the variance of the original wells, resampled with replacement and averaged. These landscapes keep the measured noise of each combination but have no dependence on composition. We repeated the decomposition on each of them and considered the contribution of order k distinguishable from noise when it exceeded the 95th percentile of the null values (Fig A).

#### Comparison with the regression models

In the HS-FR rounds, the fractions of variance carried by single-strain and pairwise terms equal the $R^2$ of the additive model and its increase when pairwise terms are added (Fig 3C). In the third lactic acid round they are close but not identical (Fig 4C), because the two plate positions of each combination are pooled here and kept as separate observations in the regression. In the first two HS-FR rounds, terms of three or more strains were within experimental noise, except third-order terms in the first round, which lay just above the null (1.6% of the variance against a threshold of 1.4%). In the third round of both functions, terms of every order exceeded noise, and those of three or more strains together accounted for 41% of the variance in HS-FR and 17% in lactic acid (Fig A).

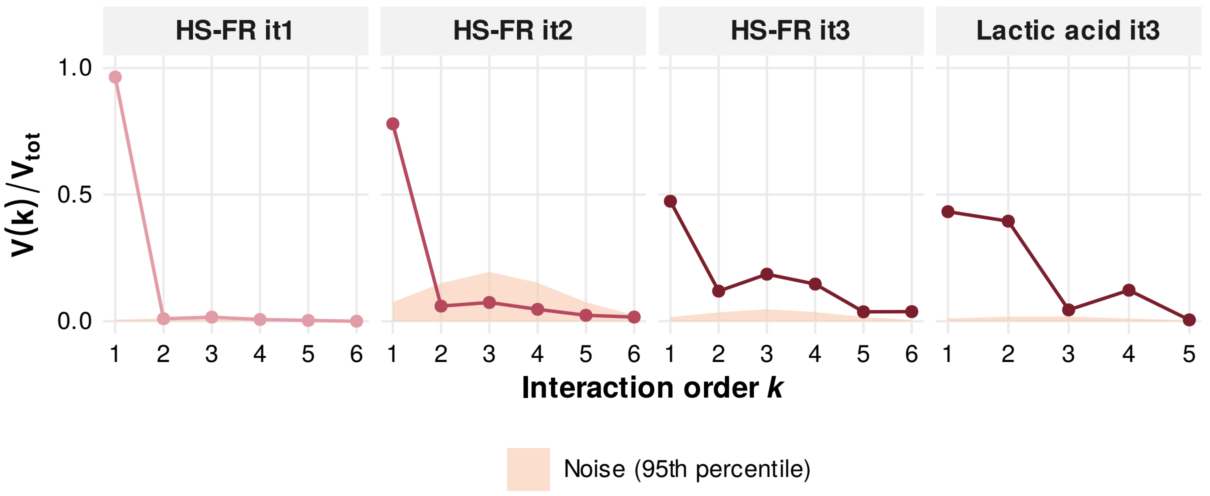


**Fig A.** Contribution of each interaction order compared with experimental noise. Fraction of the variance of the community-function landscape explained by Walsh–Hadamard terms of each order $k$, $V(k)/V_{tot}$, in the rounds with complete landscapes (HS-FR rounds 1–3 and lactic acid round 3). Points give the observed fraction, coloured by round, and the shaded band reaches the 95th percentile of the same fraction in 1000 noise-only landscapes.
